# Long-reads are revolutionizing 20 years of insect genome sequencing

**DOI:** 10.1101/2021.02.14.431146

**Authors:** Scott Hotaling, John S. Sproul, Jacqueline Heckenhauer, Ashlyn Powell, Amanda M. Larracuente, Steffen U. Pauls, Joanna L. Kelley, Paul B. Frandsen

**Author notes:** **Authors for Correspondence:** Scott Hotaling, School of Biological Sciences, Washington State University, Pullman, WA, 99164, USA, Paul B. Frandsen, Department of Plant and Wildlife Sciences, Brigham Young University, Provo, UT, 84602, USA.

## Abstract

The first insect genome (*Drosophila melanogaster*) was published two decades ago. Today, nuclear genome assemblies are available for a staggering 601 insect species representing 20 orders. In this study, we analyzed the most-contiguous assembly for each species and provide a “state of the field” perspective, emphasizing taxonomic representation, assembly quality, gene completeness, and sequencing technologies. Relative to species richness, genomic efforts have been biased towards four orders (Diptera, Hymenoptera, Collembola, and Phasmatodea), Coleoptera are underrepresented, and 11 orders still lack a publicly available genome assembly. The average insect genome assembly is 439.2 megabases in length with 87.5% of single-copy benchmarking genes intact. Most notable has been the impact of long-read sequencing; assemblies that incorporate long-reads are ~48x more contiguous than those that do not. We offer four recommendations as we collectively continue building insect genome resources: (1) seek better integration between independent research groups and consortia, (2) balance future sampling between filling taxonomic gaps and generating data for targeted questions, (3) take advantage of long read sequencing technologies, and (4) expand and improve gene annotations.

**Significance statement:** Since the first insect genome was sequenced ~20 years ago, sequencing technologies and the availability of insect genome assemblies have both advanced dramatically. In this study, we curated, analyzed, and summarized the field of insect genomics in terms of taxonomic representation, assembly quality, gene completeness, and sequencing technology. We show that 601 insect species have genome assemblies available, with some groups heavily overrepresented (e.g., Diptera) relative to others (e.g., Coleoptera). The major takeaway of our study is that genome assemblies produced with long reads are ~48x more contiguous than short read assemblies.

## Body

Since the publication of the *Drosophila melanogaster* genome (Adams, et al. 2000) sequencing and analytical technologies have developed rapidly, bringing the power of genome sequencing to an ever-expanding pool of researchers. More than 600 insects have now had their nuclear genome sequenced and made publicly available in the GenBank repository (Sayers, et al. 2021). While representing just 0.06% of the ~1 million described insects (Stork 2018), this breadth of insect genome sequencing still spans ~480 million years of evolution (Misof, et al. 2014) and roughly two orders of genome size from the tiny 99 megabase (Mb) genome of *Belgica antarctica* (Kelley, et al. 2014) to the massive genome of *Locusta migratoria* at 6.5 gigabases (Gb; Wang, et al. 2014).

Accumulating genomic resources have transformed biological research and precipitated major advances in our understanding of the origins of biodiversity (Hug, et al. 2016; McGee, et al. 2020; McKenna, et al. 2019; Seehausen, et al. 2014). Considerable progress has been driven by large-scale consortia [e.g., Human Genome Project (Collins, et al. 2003); Vertebrate Genome Project (Rhie, et al. 2021)] and for insects, the most prominent consortium has been the i5K initiative to sequence genomes for 5,000 different arthropods (i5K Consortium 2013; Robinson, et al. 2011). The rise of long-read sequencing technologies—primarily Oxford Nanopore and Pacific Biosciences (PacBio)—have also changed the landscape of genome sequencing by providing an economical means for high-throughput generation of reads that are commonly 25 kilobases (Kb) or longer (Amarasinghe, et al. 2020), thereby greatly increasing the average size of sequences used in assemblies. Genome sequencing efforts in insects, however, have not been spread evenly. Aquatic insects, as a group, are underrepresented relative to their terrestrial counterparts (Hotaling, et al. 2020). And, some orders (e.g., Diptera) are represented by far more genome assemblies than their species diversity alone would warrant—likely reflecting the model organisms within them—while many orders still have no genomic representation.

Here, we curated and analyzed the best available genome assembly for 601 insects (species or subspecies). We provide a “state of the field” perspective emphasizing taxonomic representation, assembly quality, gene completeness, and sequencing technology. We focused on taxonomic breadth rather than within-group efforts [e.g., The *Anopheles gambiae* 1000 Genomes Consortium (2017)] to gain a more holistic overview of the field. Following similar studies (e.g., Hotaling, et al. 2020; Misof, et al. 2014; Petersen, et al. 2019), we defined insects to include all groups within the subphylum Hexapoda. We downloaded metadata from GenBank for all nuclear hexapod genome assemblies on an order-by-order basis (Sayers, et al. 2021; accessed 2 November 2020). We culled this data set to only include the assembly with the highest contig N50 for each taxon and downloaded these assemblies for analysis. We acknowledge that this filtering approach may introduce biases towards the present-day for assemblies that have been improved over the years. Assemblies were classified as “short-read”, “long-read”, or “not provided” based on whether only short-reads (e.g., Illumina) were used, any amount of long-read sequences (e.g., PacBio) were used, or no information was provided. If an assembly used both short- and long-reads (a “hybrid” assembly), it was classified as a long-read assembly in our analysis.

To test if insect orders were under- or overrepresented in terms of genome assembly availability, we compared the observed number of taxa with assemblies to the expected number given the described diversity for a given order. We obtained totals for the number of insects described overall and for each order from previous studies (Bellinger, et al. 2020; Zhang 2011). We assessed significance between observed and expected representation with Fisher’s exact tests. To assess gene completeness, we ran “Benchmarking Universal Single Copy Orthologs” (BUSCO) v.4.1.4 (Seppey, et al. 2019) on each assembly using the 1,367 reference genes in the OrthoDB v.10 Insecta gene set (Kriventseva, et al. 2019). It should be noted that Collembola genome assemblies may have received slightly lower BUSCO scores in this analysis because non-insect hexapod genomes were not used to generate the Insecta gene set. We tested for differences in distributions of contig N50 or assembly size between short- and long-read assemblies with Welch’s T-tests. Next, using the BUSCO gene set, we tested whether longer genes were more likely to be missing or fragmented depending on sequencing technology (short- or long-read) with Spearman’s correlations. We defined BUSCO gene length as the full nucleotide sequence for the protein-coding portions of the consensus “ancestral” genes included in the OrthoDB v.10 Insecta gene set. An extended version of the methods and the scripts used for analysis are provided in the Supplementary Materials and GitHub repository (https://github.com/pbfrandsen/insect_genome_assemblies).

As of November 2020, 601 different insect species representing 20 orders had nuclear genome assemblies available in GenBank. These data were dominated by Diptera (*n* = 169 assemblies), Hymenoptera (*n* = 164), and Lepidoptera (*n* = 118; Fig. 1a). Four orders were overrepresented relative to their species diversity: Collembola, Diptera, Hymenoptera, and Phasmatodea (*P*, Fisher’s < 0.03; Fig. 1a). Coleoptera, with 387,100 described species (Zhang 2011), was significantly underrepresented (41 assemblies versus ~228 expected; *P*, Fisher’s < 0.01). Six orders were represented by only one genome assembly and 11 orders had no publicly available assembly. This lack of representation was particularly striking for Neuroptera (5,868 described species, Zhang 2011).

**Figure 1.**
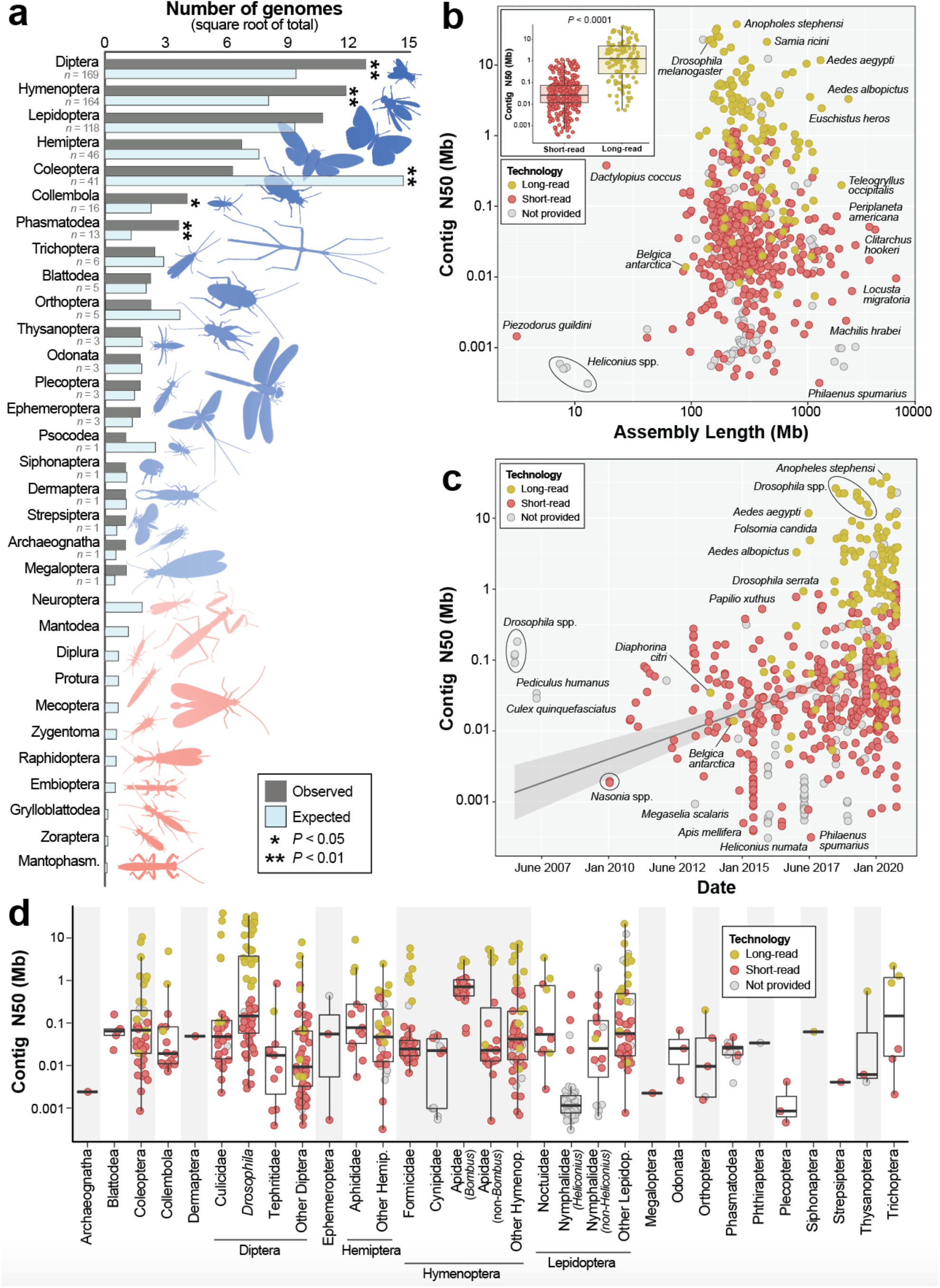
Taxonomic representation, contiguity, and the timeline of availability for the most-contiguous nuclear genome assembly for 601 insect species in GenBank as of November 2020. Only one assembly per named species or subspecies is included. (a) *The taxonomic diversity of available insect genome assemblies*. Observed versus expected numbers of genome assemblies represent the total number of available assemblies versus those that would be expected given the proportion that each order comprises of all described insect diversity. Significance was assessed with Fisher’s exact tests. One order is underrepresented (Coleoptera) while four orders are overrepresented (Diptera, Hymenoptera, Collembola, Phasmatodea). Eleven orders (light red silhouettes) have no publicly available genome assembly. A breakdown of sequencing technology by order is shown in Fig. S1. (b) *Genome contiguity versus total assembly length*. Contiguity was assessed with contig N50, the mid-point of the contig distribution where 50% of the genome is assembled into contigs of a given length or longer. The inset plot shows a comparison of contig N50 distributions for short-read (*n* = 365) versus long-read (*n* = 126) assemblies. Significance was assessed with a Welch’s T-test. A finer-scale breakdown by sequencing technology is shown in Fig. S2. (c) *The timeline of genome assembly availability for insects according to the GenBank publication date*. A steady increase in contiguity is largely precipitated by the rise of long-read sequencing. Labeled in (b) and (c): well-known or outlier genome assemblies in terms of either model status, assembly size, or contiguity. Groups of species in the same genus are labeled with black circles. (d) *Contig N50 by taxonomic group*. Generally, taxa were grouped into orders except when 10 or more assemblies were available for a lower taxonomic level (family or genus). As in (b) and (c), each point represents a single insect genome assembly.

On average, insect genome assemblies were 439.2 Mb in length (SD = 448.4 Mb; Fig. 2a) with a mean contig N50 of 1.09 Mb (SD = 4.01 Mb) and 87.5% (SD = 21%) BUSCO completeness (single and duplicated genes, combined). Substantial variation existed in all three metrics, however, with assemblies ranging from the highly incomplete assembly of *Piezodorus guildini* at just 3.2 Mb (contig N50 = 1.5 Kb, BUSCO completeness = 0.2%) to the exceptionally high-quality 140.7 Mb assembly of *D. melanogaster* (contig N50 = 22.4 Mb, BUSCO completeness = 99.9%; Fig. 2, Table S1). For orders represented by >10 taxa, Hymenoptera assemblies were the most complete (BUSCO completeness = 94%, SD = 14.3%) and Lepidoptera the least (74.6%, SD = 28.2%; Fig. 2b). At 15.3%, Lepidoptera had the lowest percentage of long-read assemblies (Fig. S1) and *Heliconius* assemblies were particularly fragmented (Fig. 1d). For families represented by >10 taxa, Drosophilidae assemblies were the most complete (BUSCO completeness = 98.4%, SD = 2%) followed closely by Apidae assemblies (97.9%, SD = 3.7%; Fig. 1d, 2b). As expected, assemblies with higher contig N50 lengths were also more complete (Fig. 2f) but assembly size had little to no effect on gene completeness (Fig. S3).

**Figure 2.**
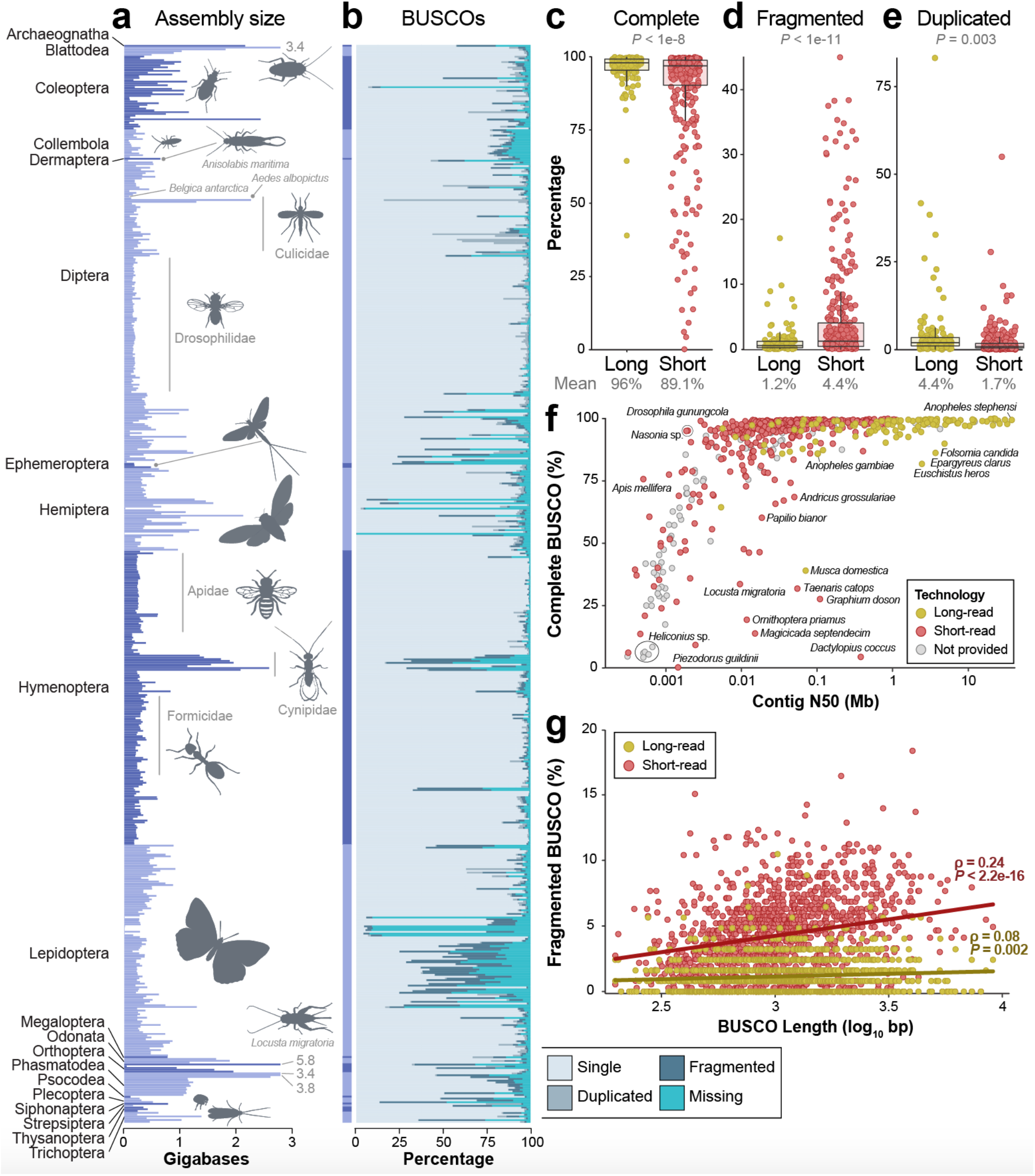
Variation in assembly size and Benchmarking Universal Single-Copy Ortholog (BUSCO) gene completeness across Insecta. (a) Assembly size for all insects, grouped by order then family. To improve visualization, the upper display limit was set to 2.8 gigabases (Gb). Four genome assemblies exceeded this value and are labeled with gray text (in Gb). Taxa silhouettes were either handmade or taken from PhyloPic (http://phylopic.org). (b) BUSCO results for each insect genome assembly. Each horizontal bar represents one assembly (*n* = 601 species) and corresponds to the same taxon in the assembly size plot to the left in (a). (c-e) Long-read versus short-read genome assembly comparisons of (c) complete BUSCOs (single and duplicated combined), (d) fragmented BUSCOs, and (e) duplicated BUSCOs only. Significance was assessed with Welch’s T-tests. (f) A comparison of BUSCO completeness versus contig N50. Each point represents the best available assembly for one taxon and groups of taxa in the same genus are labeled with black circles. Unsurprisingly, more contiguous genome assemblies also exhibit greater gene completeness. (g) Longer genes are more likely to be fragmented in insect genome assemblies, regardless of the technology used. However, a much stronger correlation exists between short-read assemblies and fragmentation of longer genes (Spearman’s ρ: 0.24, *P* < 2.2e-16) than for long-read assemblies (Spearman’s ρ: 0.08, *P* = 0.002). Unlike in (c-e), each circle in (g) represents the percent of fragmentation for that BUSCO gene across all long- or short-read assemblies. Thus, each gene is included twice (once for each technology). All 1,367 BUSCO genes in the OrthoDB v.10 Insecta gene set (Kriventseva, et al. 2019) were used except one 2.02 Kb gene that was missing in >70% of assemblies and subsequently removed from analysis and visualization. BUSCO gene lengths varied from 198 base pairs to 9.01 Kb.

The type(s) of sequence data used for genome assembly were obtained for ~82% of assemblies (long-read = 126, short-read = 365; Table S1). Long-read assemblies were more contiguous than short-read assemblies (Fig. 1b; *P*, Welch’s T-test < 0.0001), averaging contig N50 values that were ~4.4 Mb higher despite no difference in assembly size (*P*, Welch’s T-test = 0.12; Fig. S4). Gene regions were also far more complete in long-read assemblies (mean BUSCO completeness = 96%, SD = 7%) versus those generated from short-reads (89.1%, SD = 19%; *P*, Welch’s T-test < 1e-8; Fig. 2c) with 70% fewer fragmented genes (*P*, Welch’s T-test < 1e-11; Fig. 2d). Long-read assemblies, however, had ~2.6x more duplicated genes (4.4% vs. 1.7%; *P*, Welch’s T-test = 0.003; Fig. 2e). Longer BUSCO genes were also more likely to be fragmented in both short-read (Spearman’s ρ: 0.24, *P* < 2.2e-16) and long-read assemblies (Spearman’s ρ: 0.08, *P* = 0.002; Fig. 2g) but they were less likely to be missing in both when compared to shorter genes (short-read: Spearman’s ρ: −0.08, *P* = 0.002; long-read: Spearman’s ρ: −0.18, *P* = 9.7e-12; Fig. S5).

The rate at which new insect genome assemblies are becoming available is clearly accelerating (Fig. 1c). Nearly 50% (*n* = 292) of the best-available insect assemblies were accessioned in 2019–2020 (Tables S1-S2). The same period also represented a high-water mark of contiguity (mean contig N50, 2019–2020 = 1.77 Mb; Table S2). Much of the increase in contiguity was driven by long-read assemblies which rose in frequency from 0% of all assemblies in 2011–2012 to 36.1% in 2019–2020. The contiguity of long-read assemblies also sharply increased in 2017 (Fig. S6, Table S2).

We have entered a new era of insect genome biology. Since 2019, a new species has had its genome assembly deposited in GenBank every 2.3 days. These new assemblies are, on average, markedly more contiguous than those of just a few years ago. As we continue developing these resources, we offer four recommendations: first, we should recognize the community-driven nature of these data and seek better integration between research groups and consortia in terms of data sharing, best practices, and taxonomic focus. Progress towards these goals is occurring (e.g., a proposed metric system for describing genome assembly quality with associated benchmark standards from the Earth BioGenome Project, Lewin, et al. 2018) and will accelerate as more researchers integrate these standards into their own workflows. Second, new sequencing efforts should strive to balance sampling that fills taxonomic gaps and improves existing resources with targeted sampling motivated by specific questions. Both approaches are valuable and not mutually exclusive. The former—filling taxonomic gaps—is critical to broadly understanding the evolution of insects, the most diverse animal group on Earth. While the latter—targeted, question-driven sequencing—is critical to understanding specific aspects of genome biology which are often best answered using dense sampling of specific groups. Importantly, success for this recommendation will depend, in part, on our first recommendation. Better integration and communication will limit redundancy of efforts where the same species’ genome is sequenced by multiple groups simultaneously. Third, we echo the findings of the Vertebrate Genome Project (Rhie, et al. 2021)—long-read assemblies are vastly more contiguous than short-read approaches—and recommend that these technologies be embraced by insect genome scientists. And, fourth, as of 2019, only 40% of insect genome assemblies had corresponding gene annotations in GenBank (Li, et al. 2019). Expanding and refining the availability of gene annotations for insects will drive corresponding increases in the scale of taxonomic comparisons that are possible for many analyses. Overcoming this challenge of annotation quality and availability can be subdivided into two more specific calls: (1) whenever possible, annotations should be made available alongside genome assemblies in GenBank or similar public repositories and (2) researchers should consider using the NCBI Eukaryotic Genome Annotation Pipeline (Thibaud-Nissen, et al. 2016) to limit variation introduced by differing annotation approaches and maximize compatibility.

Beyond resource development, we must continue to leverage this data set to conduct new studies of insect genome biology and evolution. These efforts are beginning to emerge and are paying dividends. For instance, 76 arthropod genome assemblies were used to better understand 500 million years of evolution by characterizing changes in gene and protein content in a temporal and phylogenetic context, including the identification of novel gene families that arose during diversification with links to key adaptations including flight (Thomas, et al. 2020). Similarly, a study of 195 insect genomes revealed the high diversity of transposable elements across insects with varying levels of conservation depending upon phylogenetic position (Gilbert, et al. 2020). With genome assemblies representing 600+ taxa and ~480 million years of evolution available in a public repository, the power and promise of insect genome research has never been greater. While our focus was on insects, long-reads are likely revolutionizing genome science in virtually all taxonomic groups with untapped genomic potential existing in public repositories across the Tree of Life. The rise of long-read assemblies will, in particular, spur new understanding of previously difficult to characterize aspects of the genome (e.g., genome structure, highly repetitive regions). By continuing to build, curate, and make genomic resources publicly available, we will gain tremendous insight into genome biology and evolution at broad phylogenetic scales. We will also create a more inclusive and equitable discipline by expanding access to resources for scientists whose participation has historically been limited by financial or technological barriers.

## Acknowledgements

S.H. and J.L.K. were supported by NSF award OPP-1906015. J.H and S.U.P. were supported by the LOEWE-Centre for Translational Biodiversity Genomics which is funded by the Hessen State Ministry of Higher Education, Research and the Arts. J.S.S. was supported by an NSF Postdoctoral Research Fellowship in Biology (DBI-1811930) and an NIH General Medical Sciences award (R35GM119515) to A.M.L.

## Data availability

The data underlying this article are available in the Supplementary Materials, primarily in Table S1, with associated scripts for analysis on GitHub: https://github.com/pbfrandsen/insect_genome_assemblies.

