## Supplementary Materials for "Long-reads are revolutionizing 20 years of insect genome sequencing"

**Methods:**

*Obtaining the data*

We used the "summary genome" function in v.10.9.0 of the NCBI datasets command line tool to download metadata for all nuclear genomes for class Insecta on GenBank[1] (accessed 2 November 2020). We culled our data set to include only one genome per taxon (species or subspecies) by selecting the assembly with the highest contig N50 before downloading assemblies for analysis. We chose to use contig N50 over other metrics (e.g., scaffold N50) for two reasons: (1) to align with similar studies[2] and (2) because methods for genome scaffolding can vary dramatically and often incorporate a large number of unknown sequence (i.e., N's). Thus, while no single metric will give an ideal representation of genome assembly contiguity, in our view, contig N50 provides the most broad, comparable metric. We used TaxonKit[3] to retrieve the lineages of each species in our dataset. We first used the "name2taxid" function to retrive taxids for each species, then we used the "lineage" function to retrieve the family and order for each species.

To gather additional data for each assembly (e.g., sequencing technology), we used a custom web scraper script. Both this web scraper script and the scripts used to download and organize the metadata are available on this study's GitHub repository (https://github.com/pbfrandsen/insect_genome_assemblies). When the type of sequence data was not available in the GenBank metadata, we performed an additional literature search for each accession number on Google Scholar (https://scholar.google.com/schhp?hl=en). Using this method, we obtained the sequencing technology information for three genome assemblies (GCA_900005825.1, GCA_000004775.1, GCA_000004795.1) and have included the primary source in the "Notes" column of Table S1. We acknowledge that some genome assemblies—particularly those that were accessioned prior to 2013—are almost certain to be "short-read" based on the available technologies. But, because we had no systematic way of assigning this information (or knowing where temporal cutoffs should fall), we elected to use the objective methods described above. Our confidence in this approach was bolstered by the fact that no obvious bias appears to be present in our "no information" data set with contig N50 values spanning four orders of magnitude (see Figure S2).

*Assessing gene completeness*

To assess gene completeness, we first downloaded the assembly for each accession using the "download genome" function in v10.9.0 of the NCBI datasets command line tool. We then ran BUSCO v.4.1.4[4] in "genome mode" on each assembly using the OrthoDB v.10 Insecta gene set[5] ($n$ = 1,367 reference genes) and the "--long" option.

*Statistical analyses and visualization*

We conducted all statistical analyses and visualization in R v.3.6.1[6] with the packages *ggplot2*[7], *scales*, and *rstatix*. We performed paired T-tests with *t.test()*, ANOVA's with *aov()* followed by Tukey's HSD tests[8] with *TukeyHSD()*, and Spearman correlations by first testing if the data were normally distributed with s Shapiro's test [*shapiro.test()*] followed by a Spearman's correlation with *cor.test(method="spearman")* because none of our data were normally distributed.

**Supplementary Tables:**

See supplementary Excel file containing Tables S1-S2. Note: Table S1 can also be downloaded directly from this study's GitHub repository here: https://github.com/pbfrandsen/insect_genome_assemblies/blob/master/Table%20S1.%20GenBank%20Insect%20Genomes.xlsx

**Supplementary Figures:**

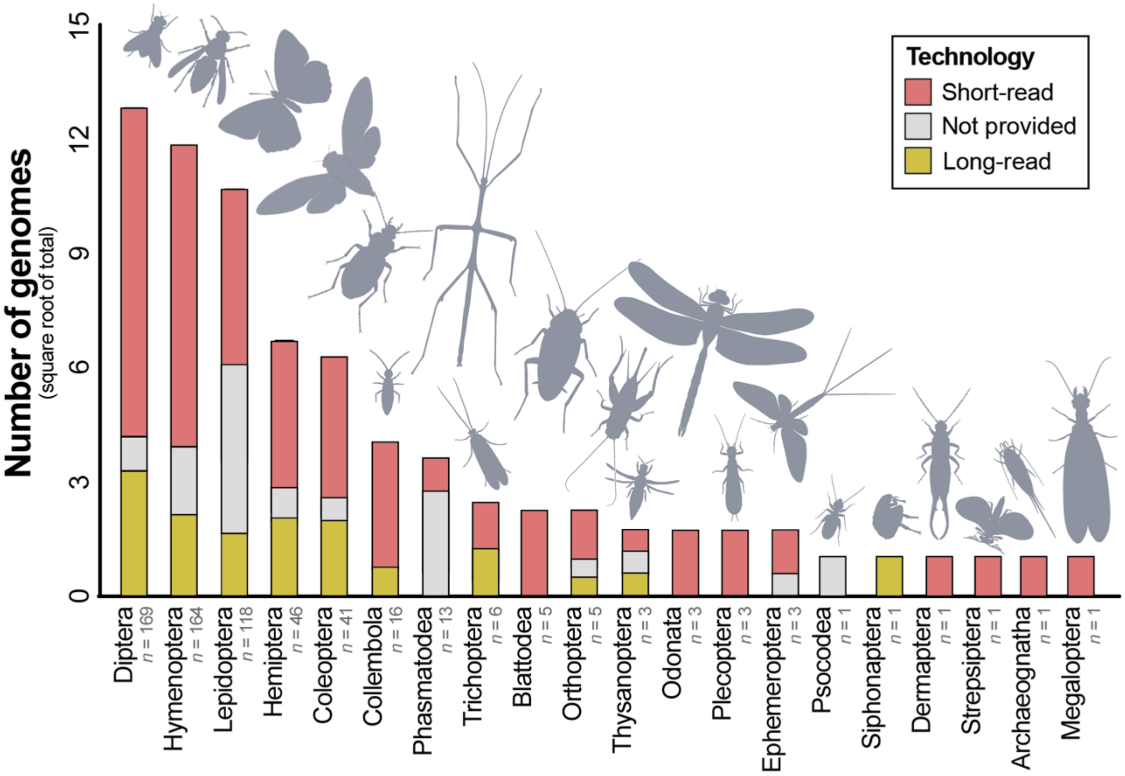


**Figure S1. The proportion of each order's genomes that have been sequenced with a given technology.** Bars align with the total number of observed assemblies on GenBank as of November 2020 and shown in Figure 1a.

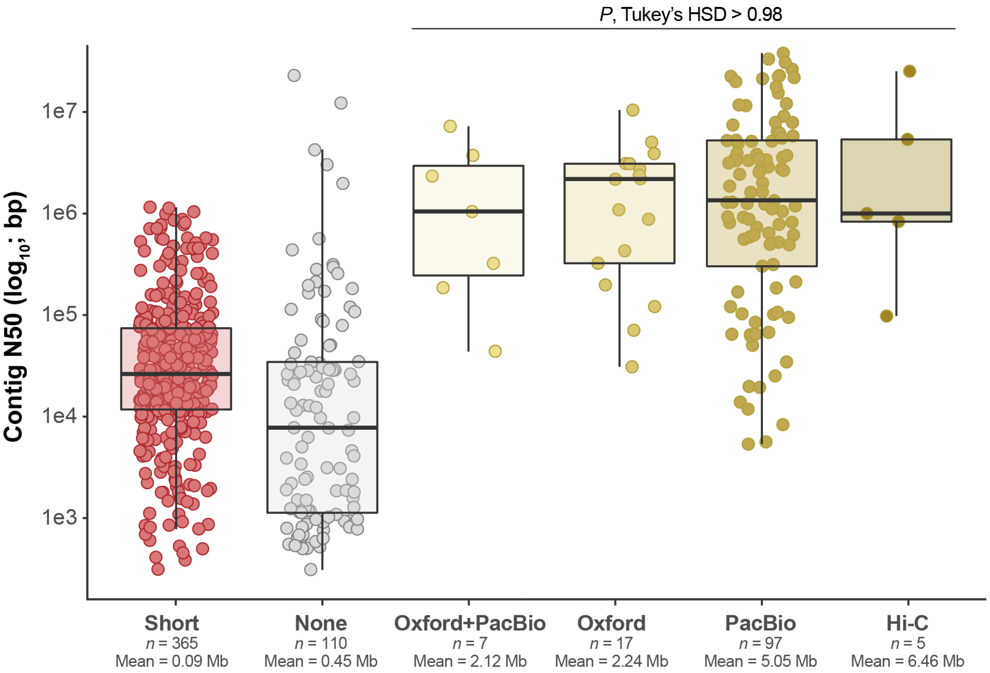


**Figure S2. The type of long-read sequence data (yellow shading) used in an assembly has little to no effect on its contiguity.** Each point represents a genome assembly for one species. Assemblies were classified based on the type of sequence data used and if proximal DNA information ("Hi-C") was used. "Short": assemblies that only incorporated sequence data with an average length < 5 kilobases (Kb). None: no information about the sequence data used were reported. "Oxford+Nanopore": incorporated both Oxford Nanopore and PacBio long-read data (with or without additional short-read data). "Oxford": incorporated Oxford Nanopore long-read data (with or without additional short-read data). "PacBio": incorporated Pacific Biosciences long-read data (with or without additional short-read data). In all cases, "Hi-C" assemblies also incorporated some type of long-read data. Significance between groups was assessed with an ANOVA followed by Tukey's HSD test.

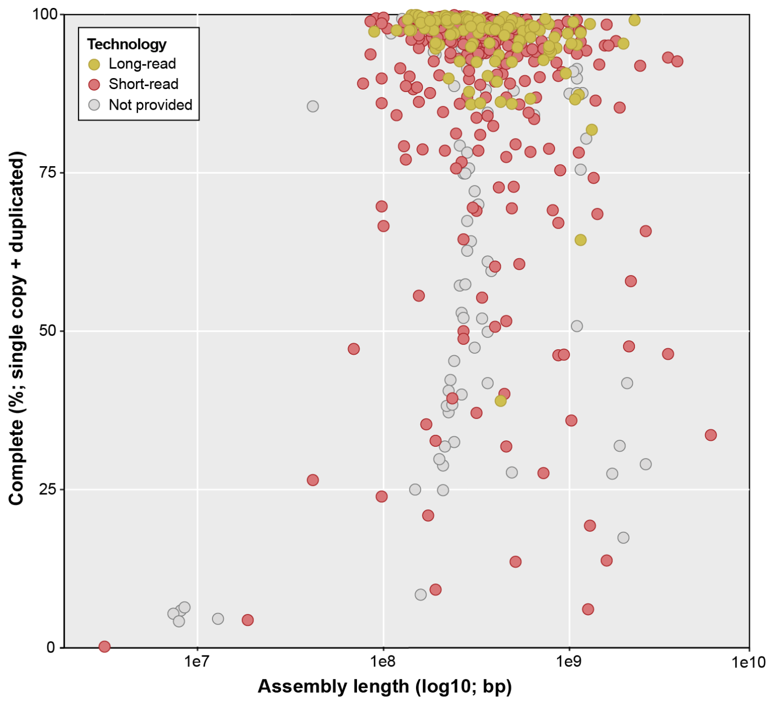


**Figure S3. Little to no relationship exists between assembly length and BUSCO completeness.** Each point represents the genome assembly for one species. Assemblies were classified based on the presence (long-read) or absence (short-read) of sequences generated with technologies that produce average read lengths greater than 5 Kb (e.g., PacBio or Oxford Nanopore).

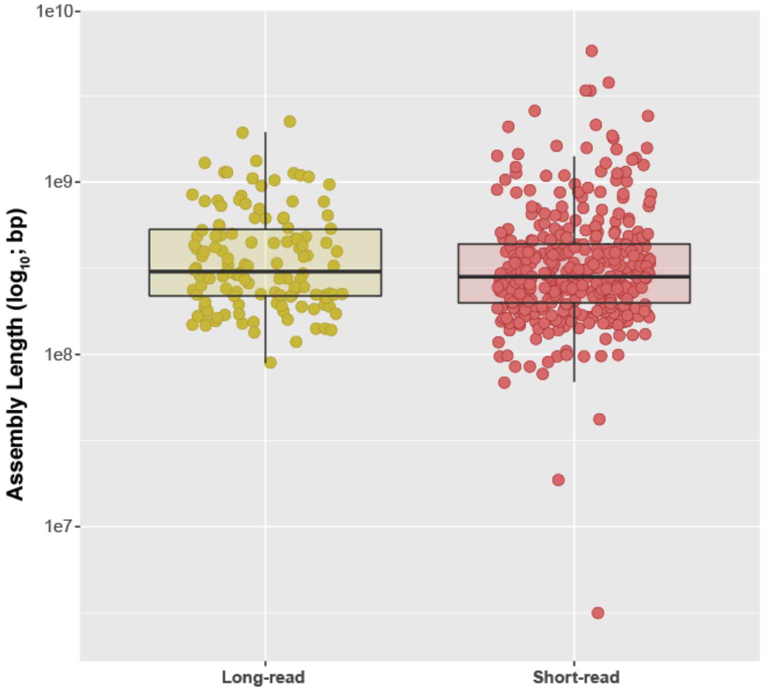


**Figure S4. There is no difference in total assembly length between long-read (*n* = 126) and short-read (*n* = 365) genome assemblies (*P,* Welch's T-test = 0.12).** Each point represents the genome assembly for one species. Assemblies were classified based on the presence (long-read) or absence (short-read) of input sequences generated with technologies that produce average read lengths greater than 5 Kb (e.g., PacBio or Oxford Nanopore).

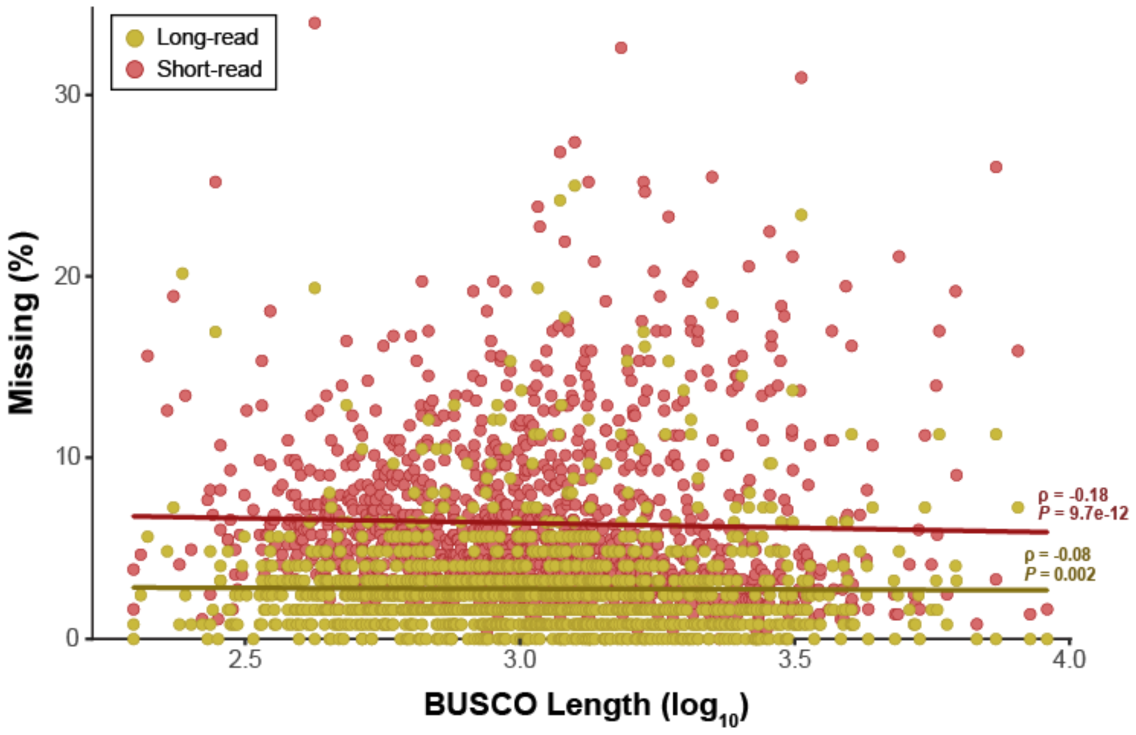


**Figure S5. Longer genes are less likely to be missing in insect genome assemblies, regardless of technology used.** However, a stronger negative correlation exists between gene length and missingness for short-read (Spearman's ρ: -0.18, *P* = 9.7e-12) versus long-read assemblies (Spearman's ρ: -0.08, *P* = 0.002). All BUSCOs included in the OrthoDB v.10 Insecta gene set (*n* = 1,367) were used with the exception of one 2.02 kilobase (Kb) gene that was missing in >70% of assemblies and was removed from the analysis and visualization. Gene lengths varied from 198 basepairs to 9.01 Kb.

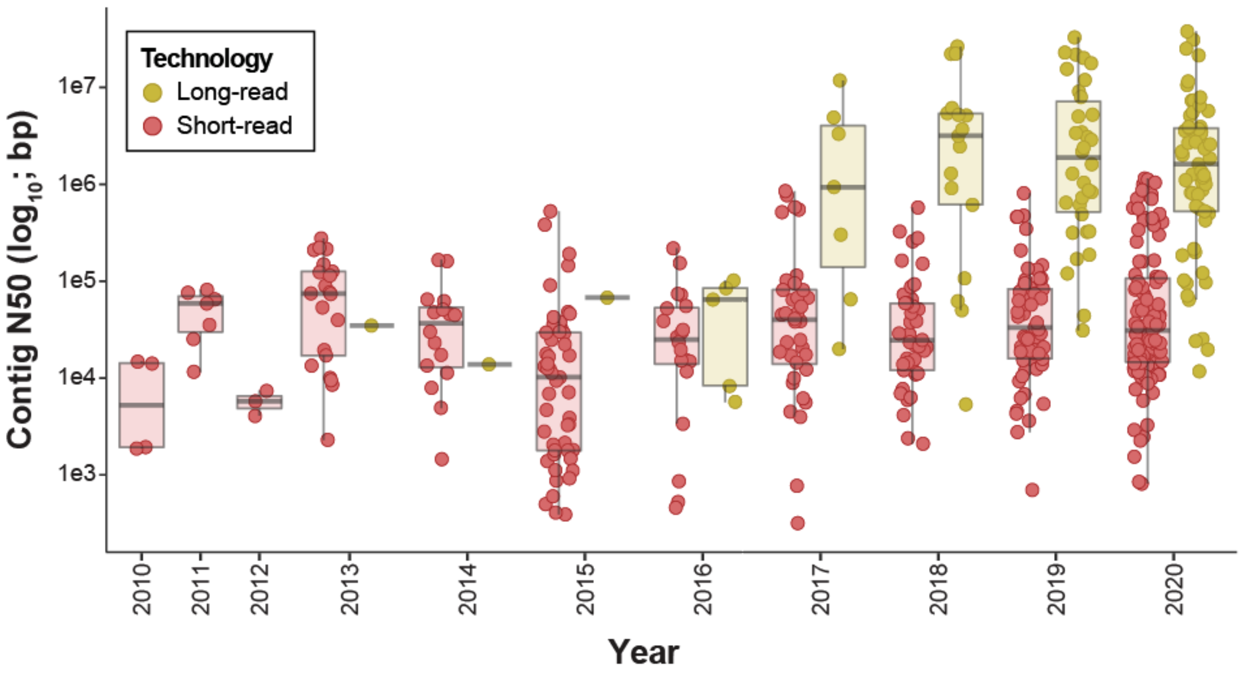


**Figure S6. Insect genome assembly contiguity by year.** While the first long-read genomes were accessioned from 2013-2016, a major shift in contiguity occurred in 2017 with long-read assemblies contiguity beginning to greatly outpace short-read assemblies. This trend has continued through 2020. Each point represents the genome assembly for one insect species. Assemblies were classified based on the presence (long-read) or absence (short-read) of sequences generated with technologies that produce average read lengths greater than 5 Kb (e.g., PacBio or Oxford Nanopore). Dark gray bars indicate median values of contig N50.
